# Secretory cells are the primary source of pIgR in small airways

**DOI:** 10.1101/2021.11.10.467794

**Authors:** Jessica B. Blackburn, Jacob A. Schaff, Sergey Gutor, Rui-Hong Du, David Nichols, Taylor Sherrill, Austin J. Gutierrez, Matthew K. Xin, Nancy Wickersham, Yong Zhang, Michael J. Holtzman, Lorraine B Ware, Nicholas E. Banovich, Jonathan A. Kropski, Timothy S. Blackwell, Bradley W. Richmond

## Abstract

**Background:** Loss of secretory immunoglobulin A (SIgA) is common in COPD small airways and likely contributes to disease progression. We hypothesized loss of SIgA results from reduced expression of pIgR, a chaperone protein needed for SIgA transcytosis, in the COPD small airway epithelium.

**Methods:** pIgR-expressing cells were defined and quantified at single-cell resolution in human airways using RNA in-situ hybridization, immunostaining, and single-cell RNA sequencing. Complementary studies in mice utilized immunostaining, primary murine tracheal epithelial cell (MTEC) culture, and transgenic mice with secretory or ciliated cell-specific knockout of pIgR. SIgA degradation by human neutrophil elastase or secreted bacterial proteases from non-typeable *Haemophilus influenzae* (NTHi) was evaluated *in vitro*.

**Results:** We found that secretory cells are the predominant cell type responsible for pIgR expression in human and murine airways. Loss of SIgA in small airways was not associated with a reduction in secretory cells but rather a reduction in pIgR protein expression despite intact *PIGR* mRNA expression. Neutrophil elastase and NTHi-secreted proteases are both capable of degrading SIgA *in vitro* and may also contribute to a deficient SIgA immunobarrier in COPD.

**Interpretation:** Loss of the SIgA immunobarrier in small airways of patients severe COPD is complex and likely results from both pIgR-dependent defects in IgA transcytosis and SIgA degradation.

**Key Messages:** <u>What is the key question</u>? Localized SIgA deficiency in small airways is an established driver of COPD pathogenesis, but the mechanism of loss remains unclear. We hypothesized loss of SIgA is due to reduced numbers of pIgR-expressing cells in SIgA-deficient small airways.

<u>What is the bottom line</u>? pIgR is primarily expressed by secretory cells in human and murine airways. Although numbers of secretory cells are similar between SIgA-deficient and SIgA-replete airways in COPD, there is reduced expression of pIgR protein, but not mRNA, in SIgA-deficient airways. Additionally, host and bacterial proteases degrade SIgA in vitro, suggesting loss of SIgA may relate to both impaired transcytosis and increased degradation.

<u>Why read on</u>? This study highlights the complexity of SIgA immunobarrier maintenance and suggests that strategies aimed at restoring the SIgA immunobarrier will need to account for both impaired transcytosis and degradation by host and/or bacterial proteases.

## Introduction

Throughout life the lungs are bombarded by inhaled particulates, microorganisms, and endogenous bacteria aspirated from the upper respiratory tract^1^. Many such irritants are deposited on the surface of small airways where bulk airflow ceases and gas transfer proceeds through diffusion^2^. To facilitate homeostasis, the small airway epithelium maintains a multifaceted defense system that protects against microbial invasion while limiting potentially injurious inflammatory responses^3^ ^4^. A key component of this system is secretory immunoglobulin A (SIgA) which lines the lumen of small airways and other mucosal surfaces^5–9^. SIgA opsonizes and agglutinates microorganisms and non-infectious irritants to facilitate their clearance through the mucociliary escalator^6^. Through a process known as immune exclusion, SIgA limits inflammation by masking pathogen-associated molecular patterns (PAMPs) and preventing them from activating inflammatory signaling cascades in the airway epithelium^6^.

In 2001, Pilette and colleagues made the seminal discovery of reduced secretory component (SC, a component of SIgA) in the airways of patients with chronic obstructive pulmonary disease (COPD)^10^, a common and often fatal lung disease associated with tobacco smoking^11^. Further, they reported that loss of SC is associated with increased neutrophilic inflammation^10^. We extended these findings to show that loss of SIgA correlates with increased bacterial invasion into small airway walls, COPD-like innate and adaptive immune activation, and small airway wall fibrosis in patients with COPD^12^ ^13^. Further, we reported that SIgA-deficient mice spontaneously develop progressive lung inflammation, small airway wall thickening, and emphysema similar to patients with COPD^14–16^, suggesting loss of SIgA immunobarrier directly contributes to COPD pathogenesis.

Regulation of SIgA on mucosal surfaces is a complex process involving multiple cell types and processes^5^. Dimeric IgA (dIgA) is made by plasma cells which reside beneath the airway epithelium in the lamina propria. To cross the airway epithelium to the luminal surface, dIgA covalently binds to the polymeric immunoglobulin receptor (pIgR) on the basolateral surface of the airway epithelium. pIgR/dIgA complexes are endocytosed and transported to the apical surface through vesicular trafficking^17^. At the apical surface an endoproteolytic cleavage event releases dIgA and the extracellular portion of pIgR called the secretory component (SC) to form SIgA. At the airway surface, SIgA is susceptible to cleavage by host and bacterial proteases^18–24^. Thus, the amount of SIgA present on the airway surface depends upon rates of production, transport, and degradation.

To better understand regulation of the SIgA barrier in the lungs, we defined cell type-specific expression of pIgR in human and murine small airways using complementary *in situ*, *in vitro*, and *in vivo* approaches and evaluated potential mechanisms accounting for an impaired SIgA barrier in small airways of patients with COPD.

## Results

### pIgR is primarily expressed by secretory cells in human airways

Secretory and ciliated cells comprise the majority of epithelial cells lining human small (< 2 mm) airways^25^. To determine which of these cell types express pIgR, we performed RNA *in situ* hybridization (RNA-ISH) for *PIGR*, *SCGB1A1* (a marker of secretory cells), and *FOXJ1* (a marker of ciliated cells) in small airways from deceased lung donors without known respiratory disease whose lungs were rejected for transplantation (e.g. control subjects).

Immunofluorescence microscopy showed areas of co-localization between *PIGR* and *SCGB1A1* but minimal overlap between *PIGR* and *FOXJ1* (**figure 1A**). Using HALO image analysis software (Indica Labs), we quantified each probe 3 μm around DAPI-stained nuclei in segmented airway epithelial cells (**online supplemental figure 1**), thus allowing us to define *FOXJ1*^+^ and *SCGB1A1*^+^ cells with minimal overlap between markers (**figure 1B**, first two panels). Using this technique, we observed higher probe expression for *PIGR* in *SCGB1A1*^+^ secretory cells compared to *FOXJ1*^+^ ciliated cells (**figure 1B**, right panel). Further, by using 2 different markers of secretory cells (*SCGB1A1* and *SCGB3A2*)^26^, we were able to account for 83% of *PIGR*^+^ cells (**figure 1C,D**) suggesting these cells are the dominant cell type responsible for *PIGR* expression in small airways. These findings were confirmed at the protein level by immunostaining in small and large airways from control patients, which showed pIgR localized in cells that contained Scgb1a1, but not FoxJ1 (**figure 1E**).

**Figure 1.**
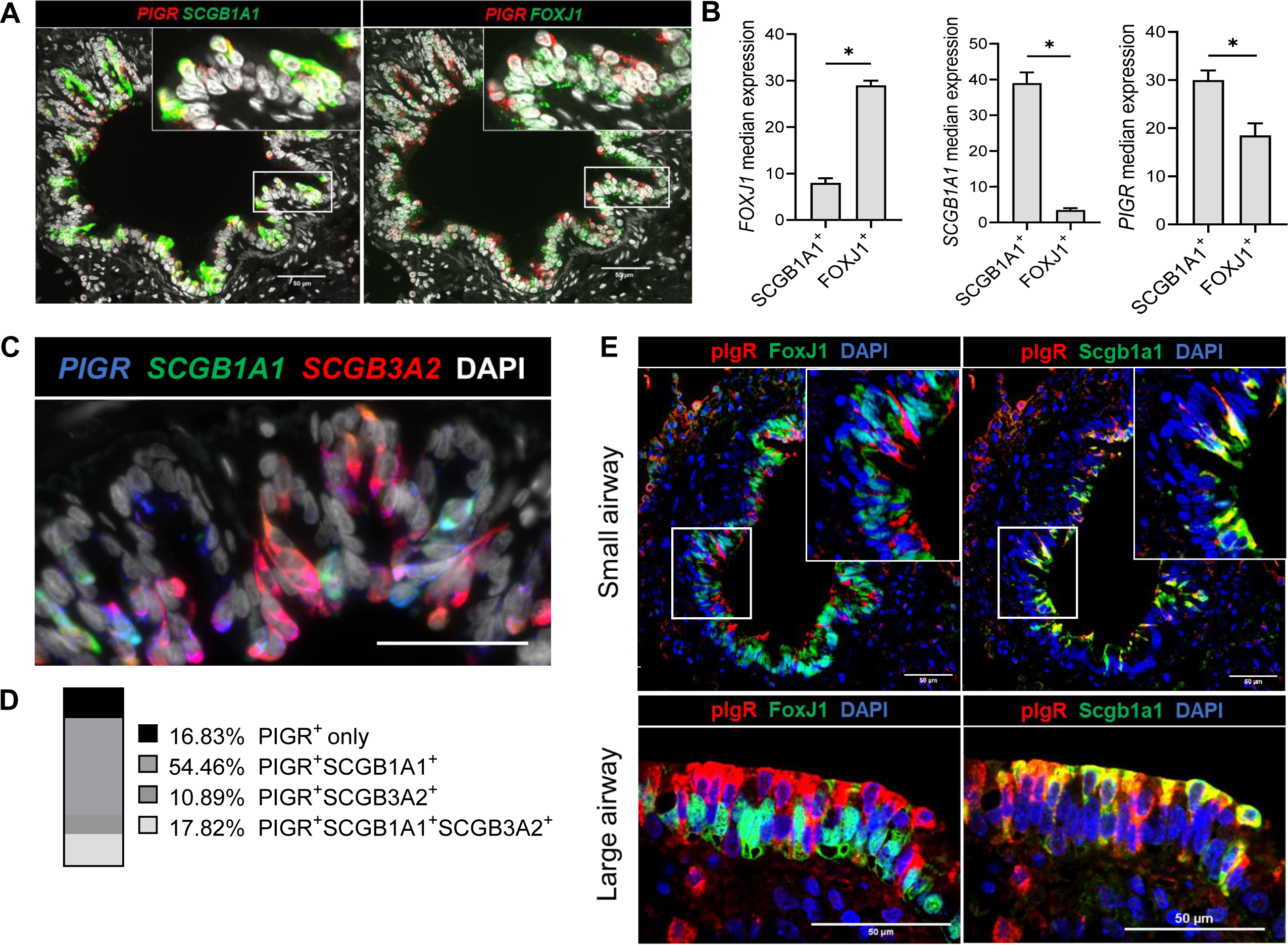
*PIGR*/pIgR is primarily expressed by secretory cells in human lung sections. (A) RNA *in situ* hybridization for *PIGR* (red), the secretory cell marker *SCGB1A1* (green, left panel), the ciliated cell marker *FOXJ1* (green, right panel), and DAPI (gray) in small airways from deceased lung donors without chronic respiratory disease. Scale bar = 50 μm. (B) *FOXJ1*, *SCGB1A1,* and *PIGR* expression (probe counts) in cells defined as *SCGB1A1*^+^ and *FOXJ1*^+^, anything under 10 probes per cell was considered background or overlap from adjacent cells. Error bars represent median values and 95% confidence interval. * = p < 0.0001 (Mann-Whitney test). (C) RNA *in situ* hybridization showing heterogenous expression of *PIGR* (blue), *SCGB1A1* (green), and *SCGB3A2* (red) and the combination of each marker in a small airway from a control patient.Scale bar = 50 μm. (D) Percentage of cells expressing *PIGR* alone or *PIGR* with secretory cell markers *SCGB1A1* and/or *SCGB3A2* (*n* = 10,908 cells (4,571 *PIGR* expressing) from 17 airways in 6 deceased lung donors without chronic respiratory disease. (E) Immunostaining for pIgR (red), Scgb1a1 (green, right panel), or FoxJ1 (green, left panel) in large and small airways from deceased lung donors without chronic respiratory disease. Scale bar = 50 μm. Inserts 2X magnification.

To gain a more comprehensive understanding of the pattern of *PIGR* expression across the respiratory epithelium, we performed single-cell RNA sequencing (scRNA-seq) of distal portions of explanted lungs from COPD patients (*n* = 11) and analyzed these samples alongside our previously published scRNA-seq data from analogous regions of lung from non-diseased controls^27^. All samples were jointly normalized and scaled prior to graph-based clustering and manual annotation using canonical markers as previously described^27^ (**figure 2A-C**, **online supplemental figure 2**). Graph-based clustering yielded three distinct populations of cells bearing markers typical of secretory cells which could be differentiated by expression of *SCGB1A1*, *SCGB3A2*, or both markers. Similar to what was observed by RNA *in situ* hybridization and immunostaining, *PIGR* expression was highest in these secretory cell populations in both COPD patients and non-diseased controls (**figure 2D**). The level of expression was similar between COPD and control samples except in ciliated cells and *SCGB1A1*^+^/*SCGB3A2*^+^ secretory cells, where expression was higher in COPD samples (**figure 2D**). Taken together, these data suggest that secretory cells are the dominant cell type responsible for pIgR expression in human airways and that loss of SIgA in COPD airways is not explained by decreased *PIGR* expression.

**Figure 2.**
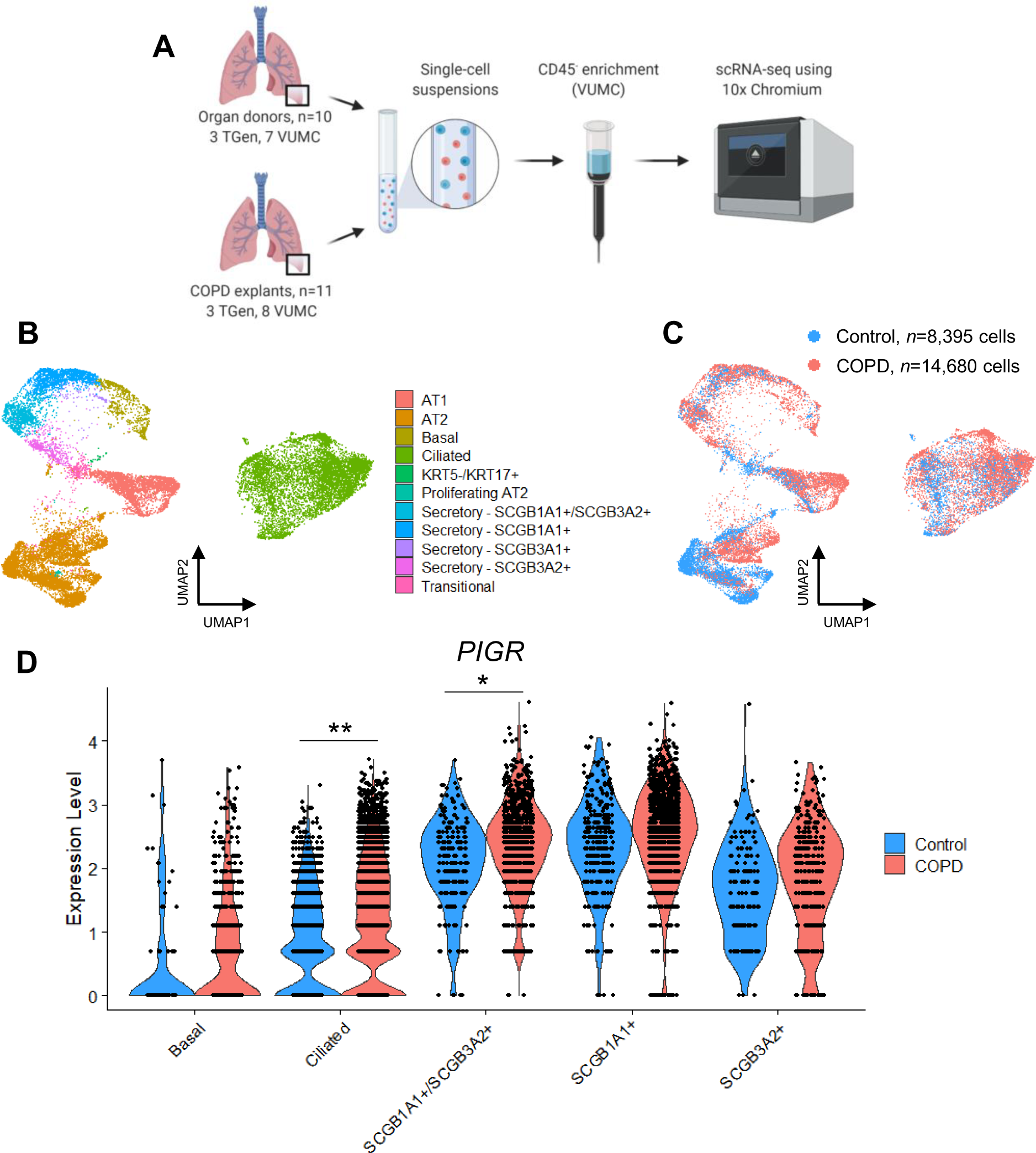
*PIGR* is primarily expressed by secretory cells in human lung tissue by sc-RNAseq. (A) Tissue processing workflow for single-cell RNA sequencing experiments. (B) UMAP-embedded image showing cell clusters derived from non-biased, graph-based clustering of EPCAM^+^ cells derived from the distal lungs of COPD and control patients. (C) UMAP-embedded showing integration of cells according to clinical status. (D) Violin plots show *PIGR* expression in COPD explants and organ donors without known chronic respiratory disease (e.g. controls). The height of each violin corresponds to the degree of expression/cell and the width of the violin corresponds to the number of expressing cells for a given level of expression. * = p < 0.001; ** = p < 0.0001 (negative binomial regression).

### pIgR is primarily expressed by secretory cells in murine airways

Similar to human airways, we found pIgR co-localized with Scgb1a1 by immunostaining but not the ciliated cell marker acetylated α-tubulin in wild-type (WT) C57Bl6/J mice (**figure 3A**). Additionally, we isolated tracheal epithelial cells from WT mice and treated them with DAP-T, an inhibitor of secretory cell maturation^28^, during terminal differentiation in air-liquid interface (ALI) culture (**figure 3A**). DAP-T markedly reduced pIgR expression in whole-cell lysates (**figure 3C,D**), consistent with the idea that secretory cells are the dominant cell type responsible for pIgR expression. Levels of acetylated α-tubulin did not differ between DAP-T and control samples, suggesting loss of pIgR was not due to impaired differentiation of ciliated cells **(figure 3C)**.

**Figure 3.**
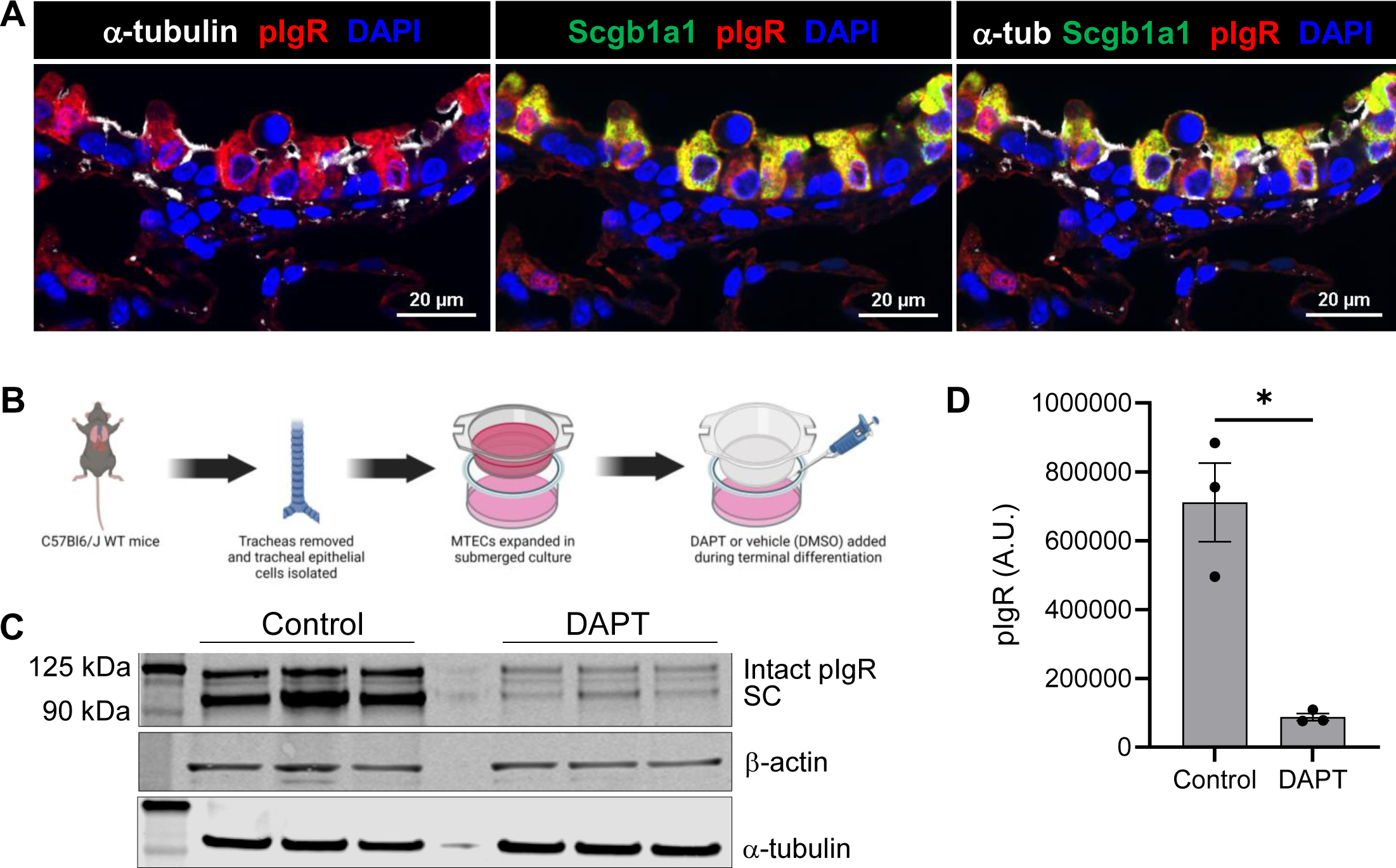
pIgR is expressed primarily by secretory cells in murine airways. (A) Immunostaining for pIgR (red), the ciliated cell marker acetylated-alpha tubulin (white), and the secretory cell marker Scgb1a1 (green) in airways from wild-type C57BL6/J mice. DAPI staining of nuclei (DNA) is shown in blue. Scale bar = 20 µm. (*B*) Diagram describing methods utilized for DAPT-T treatment in primary murine tracheal epithelial cells (MTECs). (C) Immunoblotting for pIgR in ALI-differentiated MTECs with and without addition of the gamma-secretase inhibitor DAP-T, which restricts secretory cell differentiation (*n* = 3 inserts/group). β-actin is included as a loading control and α-tubulin is included to show DAP-T did not affect ciliated cell differentiation. (D) Quantification of (C) by densitometry. All three bands of pIgR are included in the densitometry analysis. pIgR expression normalized across samples using β -actin. * = p < 0.01 (*t*-test).

Since ciliated cells outnumber secretory cells in the distal airways of humans and mice ^25^, we speculated that even low amounts of pIgR expression in ciliated cells may be relevant to SIgA transport in the distal airways *in vivo*. To investigate this further, we generated a mouse model for conditional *pigr* deletion by placing loxP sites upstream of exon 3 and downstream of exon 11 of the *pigr* gene, which includes the binding sites for dIgA (**online supplemental figure 3A**). The coding regions of green fluorescent protein (GFP) were introduced upstream and downstream of the two loxP sites, such that Cre recombinase-mediated excision of exons 3-11 results in GFP expression. pIgR^fl/fl^ mice were bred to Scgb1a1.Cre^ERT2^ mice^29^ to generate tamoxifen-inducible deletion of pIgR in secretory cells (pIgR^Δciliated^ mice) and FoxJ1.Cre mice^30^ to generate constitutive deletion of pIgR in ciliated cells (pIgR^Δciliated^ mice) (**online supplemental figure 3B**). In pIgR^Δciliated^ mice, Cre recombinase was activated by placing mice on tamoxifen chow every other week (to minimize toxicity) for 3 months (**online supplemental figure 3C**).

pIgR^Δciliated^ and pIgR^Δ^ mice were evaluated by immunofluorescence to assess pIgR expression and GFP to confirm Cre-mediated recombination. Cre^-^ littermate controls from both lines showed no expression of GFP (**figure 4A,B, left panels**). We observed weak expression of GFP in pIgR^Δciliated^ mice (**figure 4A, right panel**), suggesting low levels of *pigr* expression in FoxJ1^+^ ciliated cells. However, despite pIgR deletion from ciliated cells in these mice, many airway epithelial cells continued to express pIgR (**figure 4A, right panel, white arrows**). In contrast, we observed very few pIgR-expressing cells in pIgR^Δsecretory^ mice (**figure 4B, right panel**) and GFP expression was much more robust in pIgR^Δsecretory^ mice than pIgR^Δciliated^ mice (**online supplemental figure 4A,B**). We also performed western blotting for pIgR in whole-lung lysates from pIgR^Δciliated^ and pIgR^Δsecretory^ mice to measure global pIgR expression. In pIgR^Δciliated^ mice there was no difference in pIgR levels between Cre^+^ and Cre^-^ littermate controls (**online supplemental figure 4C,D**). In contrast, in pIgR^Δsecretory^ mice there was a marked reduction of full length pIgR in Cre^+^ mice compared to littermate control and pIgR^Δciliated^ mice.

**Figure 4.**
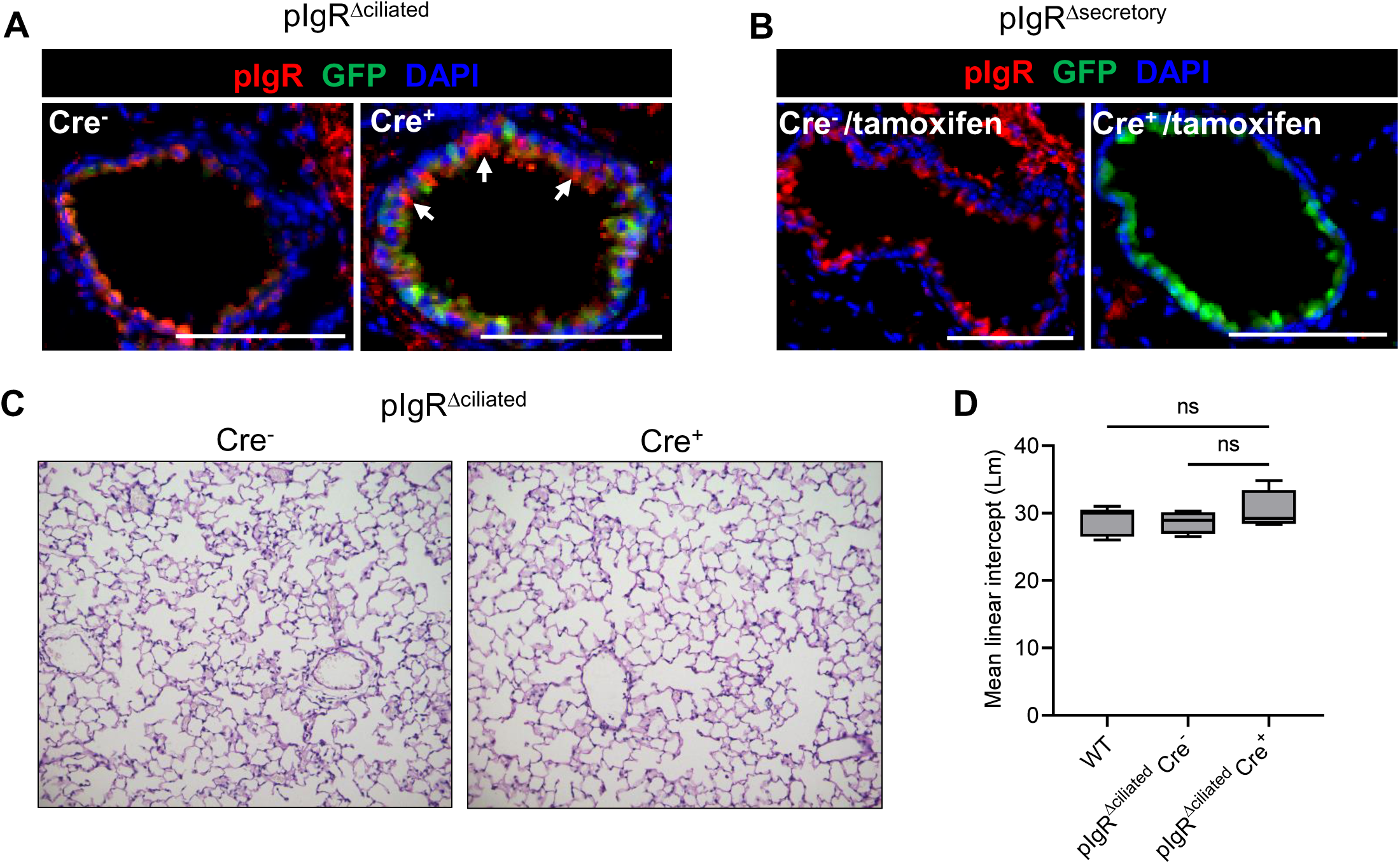
Ciliated cell-derived pIgR is dispensible for protection against emphysema. (A,B) Immunostaining for pIgR and GFP in adult pIgR^Δciliated^ and pIgR^Δsecretory^ mice. White arrows in (A) highlight pIgR^+^GFP^-^ cells in pIgR^Δsecretory^ mice. Scale bars = 50µm. (C) Low-magnification images (10x) of the lung parenchyma in 6-month-old Cre^-^ and Cre^+^ pIgR^Δciliated^ mice shows absence of emphysema in both strains (H&E). (D) Quantification of emphysema (by mean linear intercept) in 6-month-old C57Bl6 WT and Cre^+^ and Cre^-^ pIgR^Δciliated^ mice (*n*= 4-7 mice/group). Box-and-whisker plots represent median, interquartile range, and range. Results are not significant (ANOVA).

We previously noted mice with a global loss of pIgR (*pigr^-/-^* mice) lack SIgA in small airways and spontaneously develop emphysema by 6 months of age^14^. To evaluate the importance of ciliated cell-derived *pigr* for prevention of emphysema we measured mean linear intercept (Lm, a morphometric measure of emphysema) in 6-month-old pIgR^Δsecretory^ mice, Cre littermate controls, and syngeneic WT mice. We found no difference in Lm between these groups (**figure 4E,F**), suggesting SIgA transport by non-ciliated cell types is sufficient to maintain homeostasis in the murine lung. Taken together, these in vivo studies support the idea that secretory cells are the dominant cell type responsible for pIgR expression and thus play a unique role in maintaining the SIgA immunobarrier.

Changes in secretory cell numbers do not explain loss of SIgA in COPD airways.

Having established secretory cells as the dominant cell type responsible for pIgR expression, we questioned whether loss of these cells contributes to reduced SIgA in COPD airways. Using RNA-ISH, we quantified *SCGB1A1*^+^ and *SCGB3A2*^+^ cells in small airways and found a similar proportion of secretory cells between COPD and control subjects, although there was a shift towards reduced numbers of *SCGB3A2*^+^ secretory cells (**figure 5A**). To directly test whether loss of the SIgA immunobarrier was associated with a reduction in secretory cells, we performed immunostaining for SIgA as previously described ^13^ in lung sections from patients with advanced COPD and binned airways as SIgA^+^ or SIgA^-^ based on visual assessment by a pathologist (**figure 5B**). We then quantified secretory cells by RNA-ISH using probes for *SCGB1A1* and *SCGB3A2* and compared the percentage of these cell types in IgA*^+^* and IgA^-^ airways. Consistent with prior data^13^, the majority of small airways analyzed from this cohort of patients with severe COPD were SIgA-deficient (60%). However, we found no difference in numbers of *SCGB1A1*^+^ or *SCGB3A2*^+^ cells between IgA^+^ and IgA^-^ airways, though there was a trend toward reduced *SCGB3A2^+^* cells in IgA^-^ airways (**figure 5C**). These data indicate loss of secretory cells does not account for SIgA deficiency in small airways of patients with advanced COPD.

**Figure 5.**
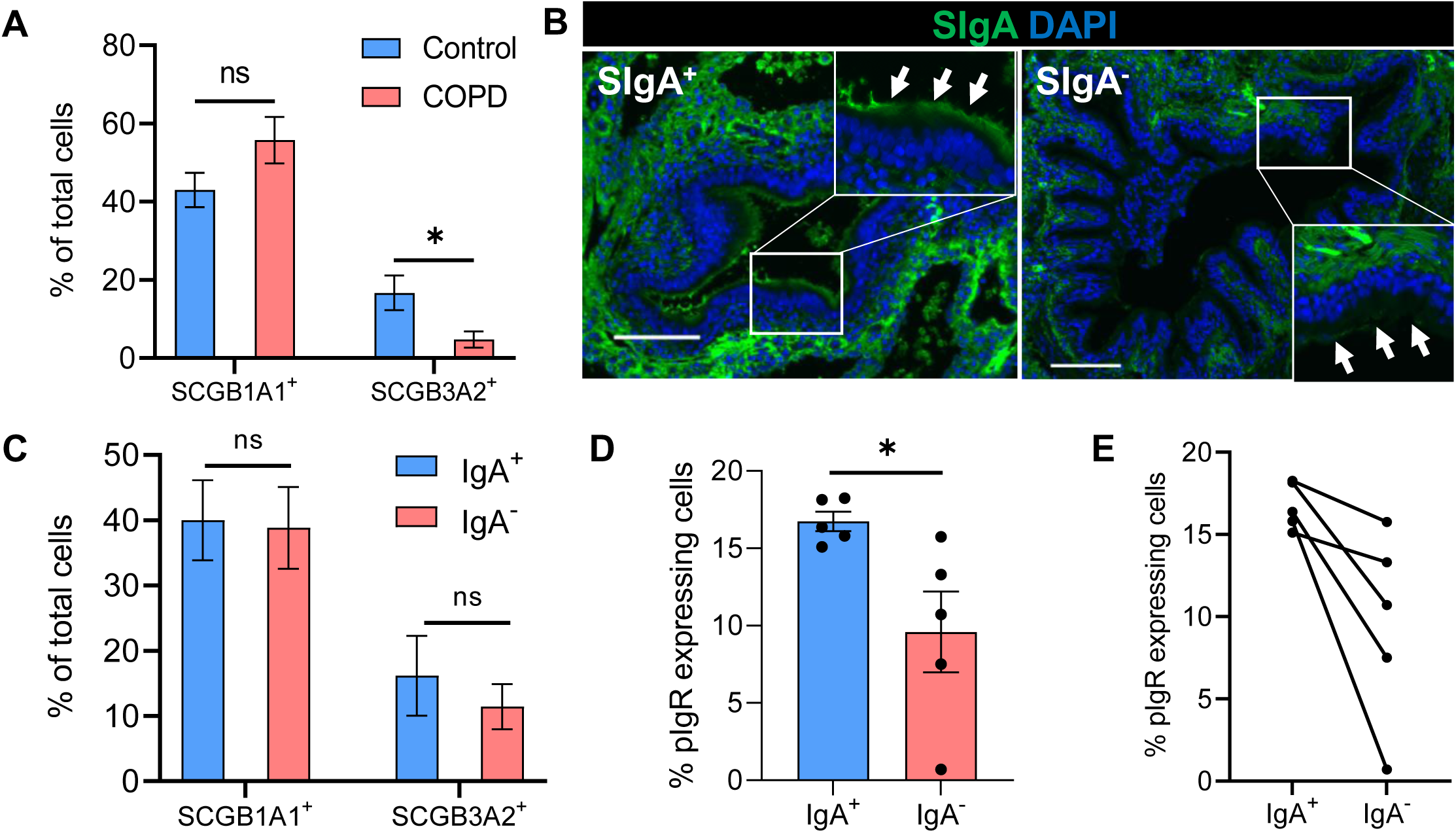
SIgA^-^ airways have reduced pIgR expression compared to SIgA^+^ airways in patients with advanced COPD. (A) Percentage of *SCGB1A1*^+^ or *SCGB3A2*^+^ cells per airway among DAPI^+^ cells as assessed by RNA-ISH in control lungs (*n* = 6, 17 airways) vs COPD (*n* = 6, 18 airways) ns = p > 0.05, * = p < 0.05, Mann-Whitney Test. (*B*) Representative images of a SIgA^+^ and SIgA^-^ small airways (based on IgA immunostaining) from the explanted lungs of a COPD patient. White arrows point to epithelial surface. Scale bars = 100 µm , inserts 2X magnification. (C). Percentage of *SCGB1A1*^+^ or *SCGB3A2*^+^ cells per airway among DAPI^+^ cells as assessed by RNA-ISH in SIgA^+^ and SIgA^-^ airways (5 patients, 18 positive airways, 28 negative) as shown in (B). (D) Average percentage of pIgR expressing cells per patient in SIgA^+^ or SIgA^-^ airways as determined by HALO analysis. * = p < 0.05, Mann-Whitney Test. (E) Paired analysis of percent of pIgR expressing cells in SIgA-vs SIgA+ airways per patient

We next considered the possibility that there was a reduction in pIgR protein expression in IgA^-^ airways despite intact *PIGR* mRNA expression seen in our scRNA-seq data (**figure 2D**). To test this, we performed immunostaining for pIgR in IgA^+^ and IgA^-^ airways. We noted a significant decrease in the percentage of pIgR^+^ cells among total cells in IgA^-^ airways (**figure 5D**) and all subjects had a higher mean perentage of pIgR-expressing cells in IgA^+^ airways relative to IgA^-^ airways (**figure 5E**). These data indicate loss of pIgR expression in secretory cells contributes to loss of the SIgA immunobarrier in patients with advanced COPD. <u>Bacterial and host proteases degrade SIgA in vitro</u>

Although we found a significant association between loss of SIgA and percentage of pIgR-expressing cells, IgA^-^ airways from some patients had a similar percentage of pIgR-expressing cells to IgA^+^ airways (**online supplemental figure 5**). Since previous studies have indicated SIgA may be degraded by bacterial or host proteases^18–24^, we speculated this might account for loss of SIgA in airways with a normal percentage of pIgR-expressing cells. To explore this further, we evaluated whether SIgA is degraded *in vitro* by secreted proteases from non-typeable *Haemophilus influenzae* (NTHi), the most common bacterium isolated from the lungs of COPD patients^31^, or sputum-derived human neutrophil elastase (HNE) which is produced by neutrophils known to accumulate around COPD small airways^13^ ^32^. We incubated SIgA derived from human colostrum with conditioned media from NTHi 1479 (a clinical isolate from a COPD patient) or with HNE and performed western blots for the IgA heavy chain and the secretory component. We found that conditioned media from NTHi cultures cleaved the IgA heavy chain but not secretory component of SIgA, consistent with IgA protease activity (**figure 6A-C**). In contrast, HNE degraded the IgA heavy chain and secretory component, forming several lower molecular weight degratory products (**figure 6D-F**). These *in vitro* data support the idea that bacterial or host proteases may also contribute to loss of SIgA in COPD.

**Figure 6.**
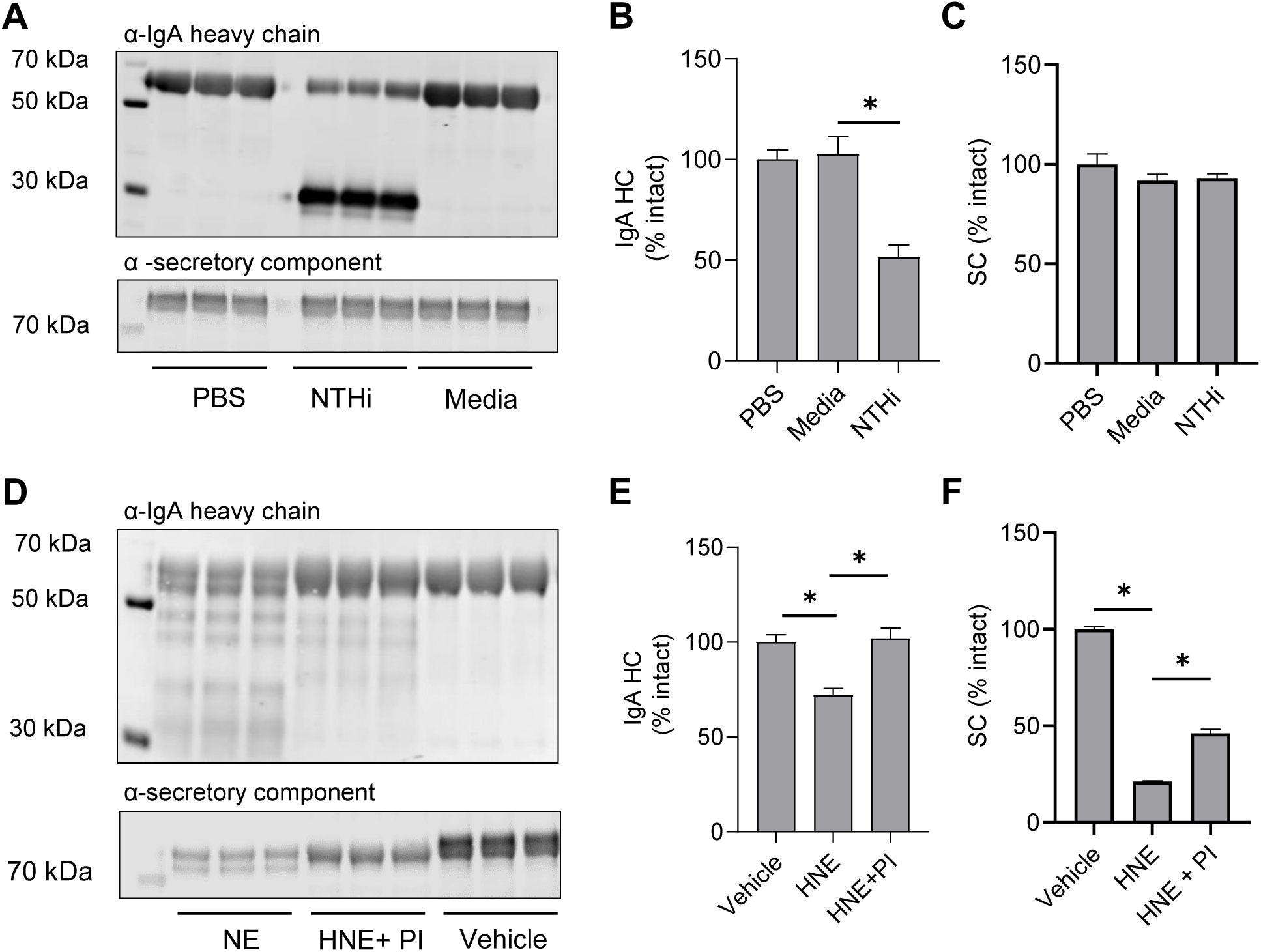
Human SIgA is degraded by bacterial and host proteases *in vitro*. (A) SIgA was incubated with conditioned media from NTHi strain 1479 for 24 hours then assessed for degradation/clevage products in the IgA heavy chain (top panel) or in the secretory component (bottom panel). PBS and non-inoculated media were used as negative controls. (B) Quantification of full length IgA heavy chain from the top panel of A. (C) Quantification of full length secretory component from the bottom panel of A. (D) SIgA was incubated with human neutrophil elastase (HNE) for 24 hours then assessed for degradation/cleavage products in the IgA heavy chain (top panel) or in the secretory component (bottom panel). The buffer used to dissolve the HNE (vehicle) was used as a negative control. A non-specific mammalian protease inhibitor was also incubated with SIgA and HNE as a control. (E) Quantification of full length IgA heavy chain from the top panel of D. (F) Quantification of full length secretory component from the bottom panel of D. * = p < 0.01 (*t*-test).

## Discussion

Loss of SIgA is common in COPD small airways and is an established driver of COPD progression in animal models^10^ ^12–16^. However, the mechanism of SIgA remains poorly understood, particularly in small airways. Here we show that secretory cells are the predominate source of pIgR in human and murine small airways and thus define a new niche for these cells in maintenance of the SIgA immunobarrier. Further, we show that loss of SIgA is associated with reduced numbers of pIgR-expressing cells despite intact *PIGR* mRNA expression. Finally, we show that bacterial and host proteases known to be present in COPD can degrade SIgA, which may also contribute to SIgA deficiency in COPD small airways.

The finding that secretory cells are the predominant source of pIgR expression is consistent with published human and murine scRNA-seq datasets^27^ ^33–37^ and with the known role of secretory cells in immune defense and airway homeostasis^35^. Although ciliated cells also produce low levels of pIgR and are more numerous than secretory cells in distal airways, we did not observe lung pathology in mice with ciliated cell-specific deletion of pIgR. This finding suggests that secretory cells play a dominant role in maintaining the SIgA immunobarrier. Interesingly, we found a shift towards reduced *SCGB3A2*^+^ secretory cells in COPD airways consistent with other studies showing a reduction in *SCGB3A2* gene expression in COPD^38^ ^39^. *SCGB3A2*^+^ secretory cells are localized predominantly in terminal and respiratory bronchioles^26^ where much of the pathology of COPD originates. The mechanism of *SCGB3A2*^+^ cell loss in COPD airways merits additional investigation.

Here we found that loss of SIgA was associated with a reduction in cells containing pIgR protein in small airways from patients with advanced COPD. Interestingly, this block appears to be at the translational or post-translational level, as we observed similar or increased levels of *PIGR* mRNA expression between COPD and control samples across cell types in our scRNA-seq dataset. Divergence between *PIGR* mRNA expression and protein levels was also described by Gohy and colleagues in large airway biopsies^40^. Since *PIGR* expression is known to be upregulated by inflammatory cytokines including IFN-γ, TNF, IL-1β, and IL-4^41−45^, persistent inflammation in small airways of COPD patients could explain our observations regarding *PIGR* mRNA expression in COPD small airways. However, the mechanism responsible for reduced pIgR protein levels in COPD remains incompletely understood and is an important area for future study.

We found that the IgA heavy chain was degraded by NTHi 1479, a clinical isolate of the most common bacterium isolated from the lungs of COPD patients during exacerbations^31^. A variety of bacteria have been shown to degrade SIgA^18^ ^19 22–24^ and it is conceivable that SIgA-degrading bacteria are overrepresented in COPD patients during stable and/or exacerbated periods^46^ ^47^. Additionally, we found that HNE is capable of degrading the IgA heavy chain and secretory component. Thus, degradation of SIgA by host and/or bacterial proteases could contribute to loss of SIgA in COPD small airways. Further, we previously showed that loss of the SIgA immunobarrier is associated with increased numbers of neutrophils within individual airways^13^ and neutrophils are spontaneously recruited to SIgA-deficient airways in mice due to chronic bacterial invasion^14^ ^15^. Thus, it is tempting to speculate that once loss of SIgA is established in an airway due to loss of pIgR, chronic bacterial invasion and neutrophilic inflammation further reduce SIgA levels in the airway in a feed-forward mechanism. Future studies examining SIgA fragmentation patterns in bronchoalveolar lavage fluid samples could prove useful for determining the specific proteases degrading SIgA.

There are several important limitations to our study. Many patients in our scRNA-seq control cohort were light smokers whereas all patients in the COPD cohort were former smokers which could have affected pIgR’s protein abundance or mRNA expression. Gohy and colleagues previously noted increased *PIGR* expression in the large airways of smokers^40^. Conversely, McGrath and colleagues found that cigarette smoke attenuated pIgR protein expression *in vivo* in response to challenge with ovalbumin and lipopolysaccharide^48^. Similarly, Rostami and colleagues showed cigarette smoke extract treatment *in vitro* reduced multiple secretory cell defense genes including *PIGR*^37^. Further studies will be required to more adequately define the impact of smoking on *PIGR*/pIgR expression. Additionally, since control samples were taken from non-diseased lung donors whose lungs were declined for transplantation, it is likely that at least some of these donors had acute lung pathology such as pulmonary hemorrhage or pneumonia which also could have influenced pIgR expression. While also subject to biases, future studies involving unaffected tissue from lung nodule/cancer resections would be complementary if matched to COPD samples by age and smoking status. Finally, our study does not exclude the possibility that there are additional pIgR-independent defects in SIgA transcytosis. Future in vitro experiments utilizing COPD-relevant models of abnormal epithelial differentiation such as that recently reported by Rao and colleagues^49^ would be helpful in examining this further.

In summary, our study suggests that secretory cells are predominantly responsible for maintaining the SIgA immunobarrier in small airways, and that loss of SIgA in COPD small airways is complex and likely due to both reduced expression of pIgR and degradation by host and bacterial proteases. Strategies to restore the SIgA immunobarrier will likely have to account for multiple layers of dysregulation in COPD.

## Funding

This work was supported by IK2BX003841 from the Department of Veterans Affairs (B.W.R.), T32 5HL094296 (J.B.B., T.S.B. PI), NIH/NHLB1 K08 HL138008 (B.W.R.), l01BX002378 from the Department of Veterans Affairs (T.S.B.), NIH/NIAID R01 AI130591 and NIH/NHLBI R35 HL145242 (M.J.H.), NIH HL126176 (L.B.W.), NIH/NHLBI R01HL145372 (J.A.K./N.E.B.), K08HL130595 (J.A.K.), and the Doris Duke Charitable Foundation (J.A.K.).

## Acknowledgements

The authors would like to thank Angela Jones and Neha Joshi in the Vanderbilt Technologies for Advanced Genomics (VANTAGE) Core for assistance with scRNA-seq. Additionally, the authors would like to thank the Vanderbilt Lung Transplant Team and the Vanderbilt Department of Surgical Pathology for assistance obtaining lung explants.

## Methods

### Human samples

The explanted lungs of COPD patients were obtained after informed consent according to Institutional Review Board (IRB)-approved protocols from Vanderbilt University Medical Center and Norton Thoracic Institute (Vanderbilt IRB #060165, 171657; Norton IRB #20181836). Control samples were obtained from deceased organ donors and were exempt from IRB review. Demographic and clinical information for human samples is provided in **Supplemental Table 1**. Demographic and clinical information for control lungs used for scRNA-seq have been published previously but are included here for reference^27^.

### Mouse models

All mouse experiments were performed according to a protocol approved by the Institutional Animal Care and Use (IACUC) Committee at Vanderbilt University Medical Center. Scgb1a1.Cre^ERT2^ mice were generated in the lab of Dr. Brigid Hogan^29^ and were obtained from the Jackson Laboratory (catalog #16225). Transgene activation was induced with tamoxifen citrate-impregnated (400 mg/kg) chow (Envigo, catalog no. 130860) as described in the main text. FoxJ1.Cre mice were originally developed in the lab of Dr. Michael Holtzman^30^ and were a gift from the developers. The flox-pIgR mouse line was developed with Ingenious Targeting Labs (Ronkonkoma, NY). A targeting vector was generated in which loxP sites were inserted upstream of *pigr* exon 3 and downstream of exon 11, which includes the binding domains for pIgR^5^. The coding regions of the GFP gene were introduced upstream and downstream of the two loxP sites, such that excision of exons 3-11 via Cre-recombinase results in GFP expression. Selection was achieved by introduction of a neomycin selection cassette flanked by FRT sites downstream of exon 11. This construct was electroporated into C57BL/6J FLP embryonic stem cells which were screened for neomycin resistance. Positive cells were injected into BALB/c blastocysts and then surgically inserted into pseudo-pregnant females. Resulting chimeras with a high percentage of black coat color were mated to C56BL/6J mice to generate germline neo-deleted mice, which was confirmed by Southern blot. A map of the targeting vector is included as **Supplemental Fig. 3**. C57BL/6J mice were purchased from the Jackson Laboratory (catalog #000664) or bred in-house after originally being purchased from the Jackson Laboratory. Approximately equal numbers of male and female mice 2 months of age or older were used in all experiments.

### Single-cell RNA sequencing

#### Tissue Processing

Biopsies from each lung sample were digested in an enzymatic cocktail (Miltenyi Multi-Tissue Dissociation Kit or collagenase I/dispase II 1 μg/ml tissue) using a gentleMACS Octo Dissociator (Miltenyi, Inc). Tissue lysates were serially filtered through sterile gauze, 100 μm, and 40 μm sterile filters (Fisher). Single cell suspensions then underwent cell sorting using serial columns (Miltenyi Microbeads, CD235a, CD45) or FACS. At VUMC, CD45^-^ and CD45^+^ populations mixed 2-3:1 were used as input for generation of scRNA-seq libraries. At TGen, Calcien-AM was used to stain live cells and 10,000-15,000 total live cells were sorted directly into the 10X reaction buffer and transferred immediately to the 10X 5’ chip A (10X Genomics).

#### scRNA-seq library preparation and next-generation sequencing

scRNA-seq libraries were generated using the 10X Chromium platform 5’ library preparation kits (10X Genomics) targeting 10,000 cells per sample. Next generation sequencing was performed on an Illumina Novaseq 6000 using Illumina S1 flow cell with paired end reads. Reads with Phred scores <30 were filtered out and 10X Genomics CellRanger v3 (for VU_COPD_29 and VU_COPD_34) or v5.0.0 (all other samples) was used to align reads to the GRCh38 reference genome.

#### scRNA-seq analysis

Post-alignment analysis was performed using the Seurat v4.0.1 package in R. Quality control filtering was performed using detected genes (minimum 750) and mitochondrial reads percent (minimum 0%, maximum 15%). All data (COPD, and control samples) were jointly normalized and scaled using the SCTransform function in Seurat^50^. Epithelial (EPCAM^+^), stromal (EPCAM^-^/PTPRC^-^) and immune (PTPRC^+^) cells were then extracted into independent objects which underwent SCT-Integration^50^ followed by principal component analysis, recursive clustering, UMAP embedding, and cell-type annotation. Clusters comprised of non-physiologic marker combinations (i.e., PTPRC^+^/EPCAM^+^ cells) were presumed doublets and excluded from further downstream analysis. Cell-type specific differentiation expression analysis was performed as previously described^27^. For this dataset, only EPCAM^+^ epithelial populations were examined. The code used to generate the dataset is available at https://www.github.com/kropskilab/copd/. Gene Expression Omnibus (GEO) repository number pending.

#### RNA *in situ* hybridization

RNA *in situ* hybridization was performed on formalin-fixed, paraffin-embedded (FFPE) tissue using the RNAscope platform (ACDBio, manual multiplex kit V2). Probes used include *PIGR*-C1, *SCGB3A2*-C2, *SCGB1A1*-C3, and *FOXJ1*-C2. Positive and negative control probes were also used to validate RNA integrity and signal specificity. TSA Plus fluorophores (fluorescein, Cy3, and Cy5) were purchased from Akoya Biosciences. Fluorophores were used at concentrations between 1:750 and 1:1500. Slides were imaged on a Nikon spinning disk confocal microscope (Yokogawa CSU-X1 spinning disk head, Andor DU-897 EMCCD, high-speed piezo [z] stage and a four-line high-power solid state laser launch) or on a Keyence BZ-710.

#### HALO analysis

Quantification of RNAscope images and pIgR immunostaining was performed using HALO image analysis software version 1-3 with the FISH-IF module (Indica Labs). Only cells within airway epithelia were quantified. DAPI was used to identify individual cells and a cell radius of 3 μm outside of the DAPI stained area was set to identify probes/signal within each cell while minimizing probe/signal spill over from adjacent cells or signal on the apical surface of the epithelium. Cells were segmented aggressively to avoid cell overlap. Minimum threshold for probe intensity for RNA-ISH or staining intensity for IF and contrast threshold was determined for each set of staining by adjusting settings for best mask fitting using real-time tuning to eliminate nonspecific signal using irrelevant areas outside of the airway epithelium (predominantly smooth muscle cells) as baseline for zero expression/staining. Probes were also segmented aggressively.

#### Immunostaining

For FFPE tissue, 5 μM sections were deparaffinized then antigen retrieval was performed using citrate buffer at pH 6. Samples were then permeabilized with 0.1% Triton prior to blocking with 1% BSA in PBS. Antibodies were incubated overnight at 4°C. Antibodies used were: pIgR (rabbit anti-human, Sigma, HPA012012, 1:1000 amplified), pIgR (goat anti-mouse, R&D systems, AF2800, 1:100 amplified), FoxJ1 (rabbit anti-human, Sigma, HPA005714, 1:500 amplified), acetylated α-tubulin (mouse anti-mammal, Santa Cruz, sc-23950, 1:100), Scgb1a1 (rat anti-human, R&D systems, MAB4218, 1:50 amplified), Scgb1a1 (rabbit anti-mouse, Abcam, ab40873, 1:1000), GFP (chicken anti-GFP, Aves labs, GFP-1010,1:250), and IgA (rabbit anti-human, Dako, A0262, 1:75). For amplified targets the appropriate HRP-conjugated secondary was then incubated at 1:250 in PBS for 45 minutes-1hr followed by TSA fluorophore treatment (Akoya). For unamplified targets appropriate Cy3 or FITC labeled secondary antibodies were used at 1:100. Images were taken on a Nikon spinning disk confocal microscope (Yokogawa CSU-X1 spinning disk head, Andor DU-897 EMCCD, high-speed piezo [z] stage and a four-line high-power solid state laser launch) or on a Keyence BZ-710.

#### SIgA assessment

Airways were binned into SIgA^+^ and SIgA^-^ based on fluorescent signal of IgA on the luminal surface as previously described^13^.

#### Western blots

Whole-tissue lysates were prepared from flash-frozen lung tissue (for Flox-pIgR experiments) or whole-cell lysates (for MTEC experiment) using RIPA buffer supplemented with protease inhibitors. Protein concentrations were determined by bicinchoninic acid (BCA) assay and 15 μg of protein (or 8 µg of SIgA for *in vitro* degradation assays) was separated on a 10% acrylamide gel and transferred to a nitrocellulose membrane. Immunoblotting was performed using antibodies against murine pIgR (goat anti-mouse, R&D catalog no. AF2800, 1:1000), β-actin (mouse anti-multispecies, Sigma A5316, 1:40,000), α-tubulin (1:1000), IgA heavy chain (rabbit anti-human, Dako, A0262 1:1000) and secretory component/pIgR (goat anti-human, R&D AF2717, 1:500) followed by detection with fluorescent secondary antibodies (1:10,000) using the Odyssey Imaging System (Li-Cor). Densitometry was performed using Image Studio, and standardized to β-actin.

#### Cell culture

*Murine tracheal epithelial cell (MTEC) isolation, culture, and DAP-T treatment*. MTECs were isolated and cultured as described by You and colleagues^51^. Briefly, adult C57BL/6J mice were euthanized and tracheas isolated and split longitudinally to expose the lumen. Tracheas were placed in 1.5 mg/ml Pronase E in Ham’s/F12 containing antibiotics and incubated overnight at 4°C. Cells were dislodged from the tracheas by gentle agitation in Ham’s/F12 containing 10% fetal bovine serum (FBS), collected by centrifugation at 500 x g, and resuspended in MTEC basic medium^51^ containing 10% FBS. Cells were then plated onto a Primaria tissue culture dish (Corning, catalog #353801) for 3–4 hr. at 37°C with 5% CO_2_ to adhere contaminating fibroblasts. Non-adherent cells were then collected and resuspended in MTEC complete media ^51^ and seeded at a density of 0.8 x 10^5^ onto Transwell inserts (Corning, catalog #3470) pre-coated with 0.05 mg/ml rat tail Type 1 collagen. After cells reached confluence (∼Day 7-10), they were lifted to generate air-liquid interface (ALI) cultures with MTEC 2% NuSerum/RA medium^51^ added only to the lower chamber. For the DAP-T experiment, 10 µM DAP-T (Abcam, catalog #120633) was added to media in the basolateral chamber for 14 days of ALI culture. Control wells were treated with diluent (DMSO) only.

#### In vitro SIgA degradation experiments

For the NTHi degradation experiment, NTHi strain 1479 (a gift from Dr. Brahm Segal, University of Buffalo) was grown overnight at 37°C on chocolate agar plates (Hardy Diagnostics). A single colony was used to inoculate 20 ml of brain-heart infusion (BHI) media (Sigma-Aldrich) supplemental with 10 μg/ml NAD and 10 μg/ml Hemin (both from Sigma-Aldrich). The culture was incubated for 4 hours at 36°C with constant shaking and then used to inoculate an additional 200 ml of liquid media. After another 4 hours of growth, bacteria were pelleted and the conditioned media filtered using a 0.22 μm PES filter. This media (25uL) was incubated with 25 μg (1ug/uL, 25uL) SIgA from human colostrum (Athens Research and Technology, 16-13-090701). For the human neutrophil elastase (HNE) experiment, 25 μg of SIgA was incubated with 10 μg sputum-derived human neutrophil elastase (1 ug/uL in 0.05 M NaOAc pH 5 containing 0.1 M NaCl, Elastin Products Company, SE563GI) with or without a protease inhibitor (Sigma, P8340 1:100).

#### Statistics and graphing

Mice were randomly assigned to study groups and all animals were included in each analysis. Results are presented as mean ± standard error of the mean unless otherwise noted. GraphPad Prism 9 was used for statistical testing and graphing. Type of test and *p* value are noted in each figure legend.

## Online Supplemental Figure Legends

**Supplemental Figure 1.**
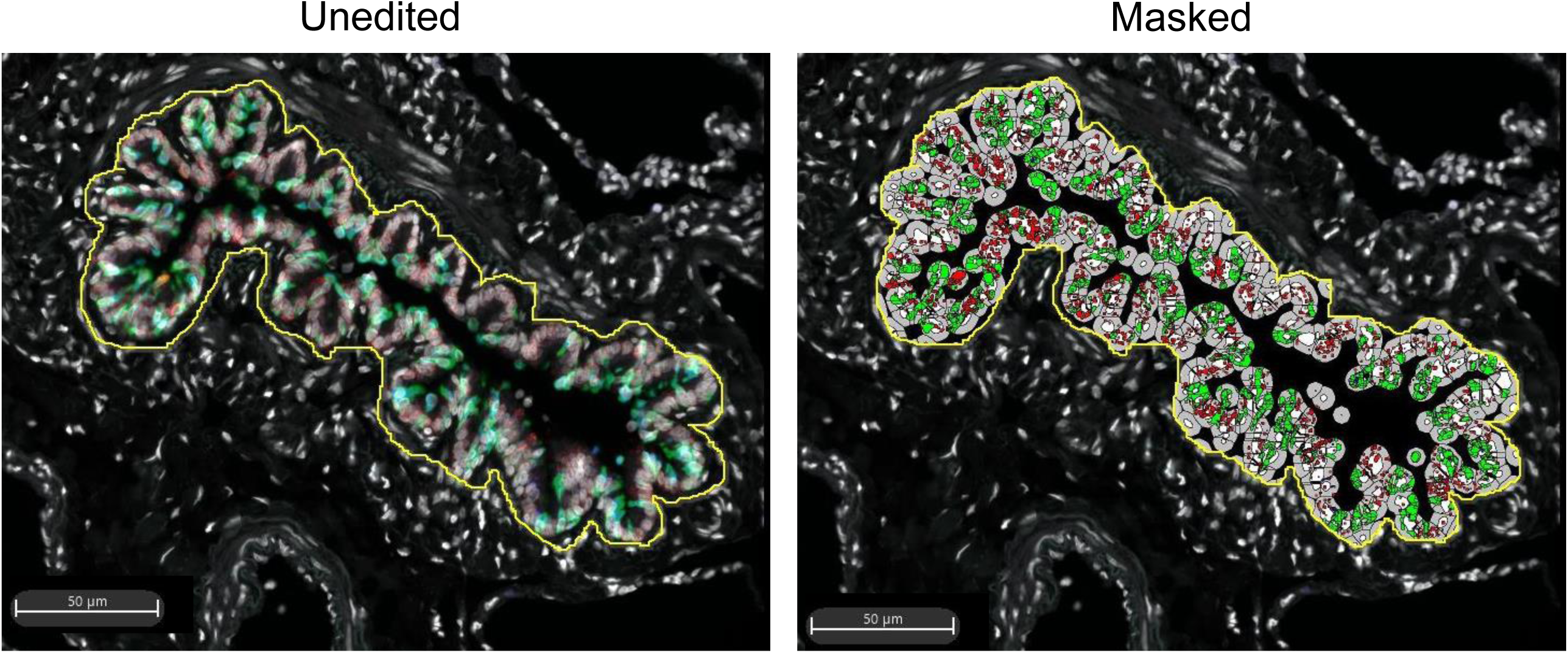
Example of region of interest definition and masking using HALO image analysis software. Images of a small airway from a non-diseased donor with and without masking performed using HALO. The region of interest was defined as shown by the yellow line and was drawn at the base of the airway epithelium.

**Supplemental Figure 2.**
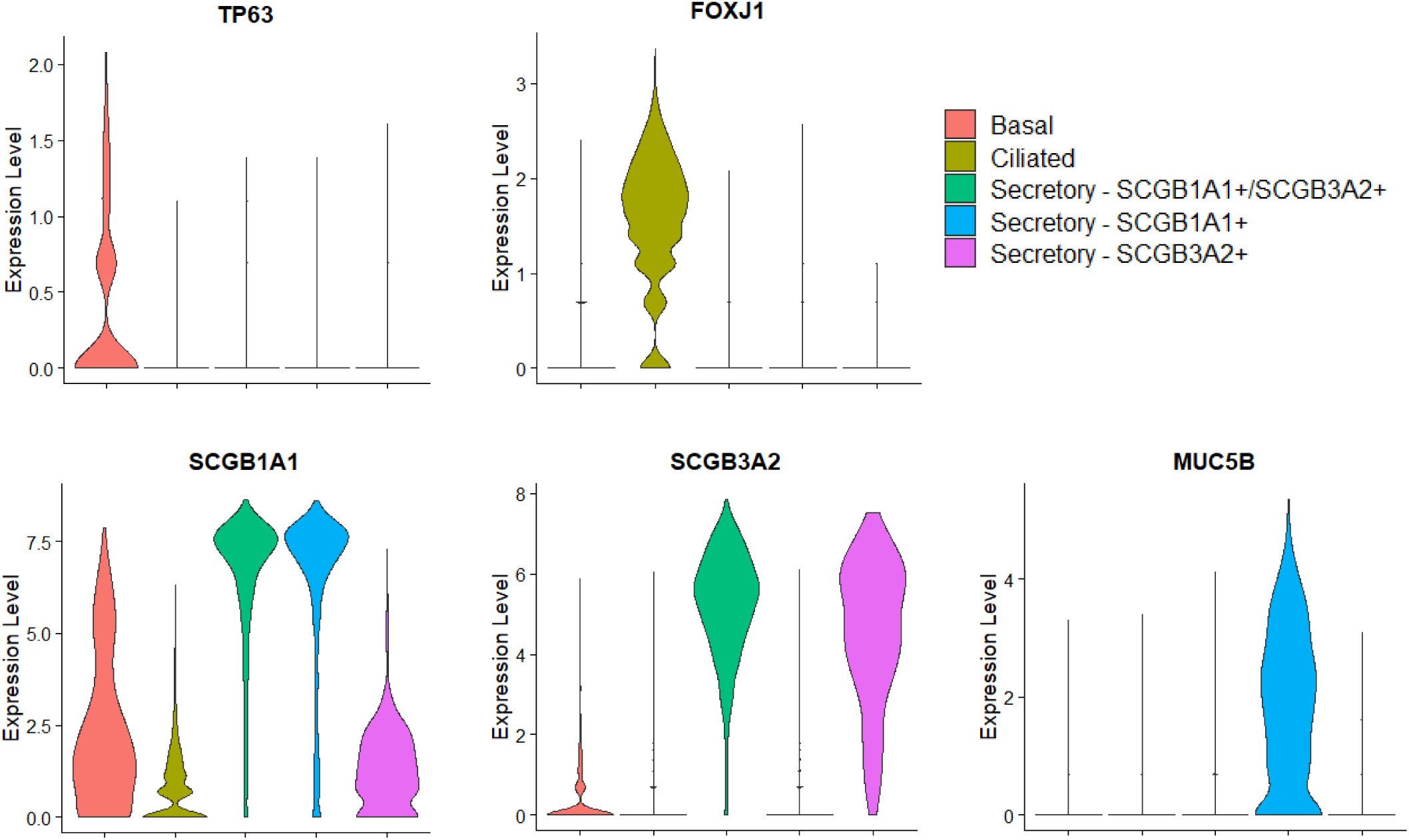
Canonical markers used to define cell types in the merged control and COPD dataset. Violin plots showing expression of markers used in manual annotation across cell types. The height of each violin corresponds to the degree of expression/cell and the width of the violin corresponds to the number of expressing cells for a given level of expression.

**Supplemental Figure 3.**
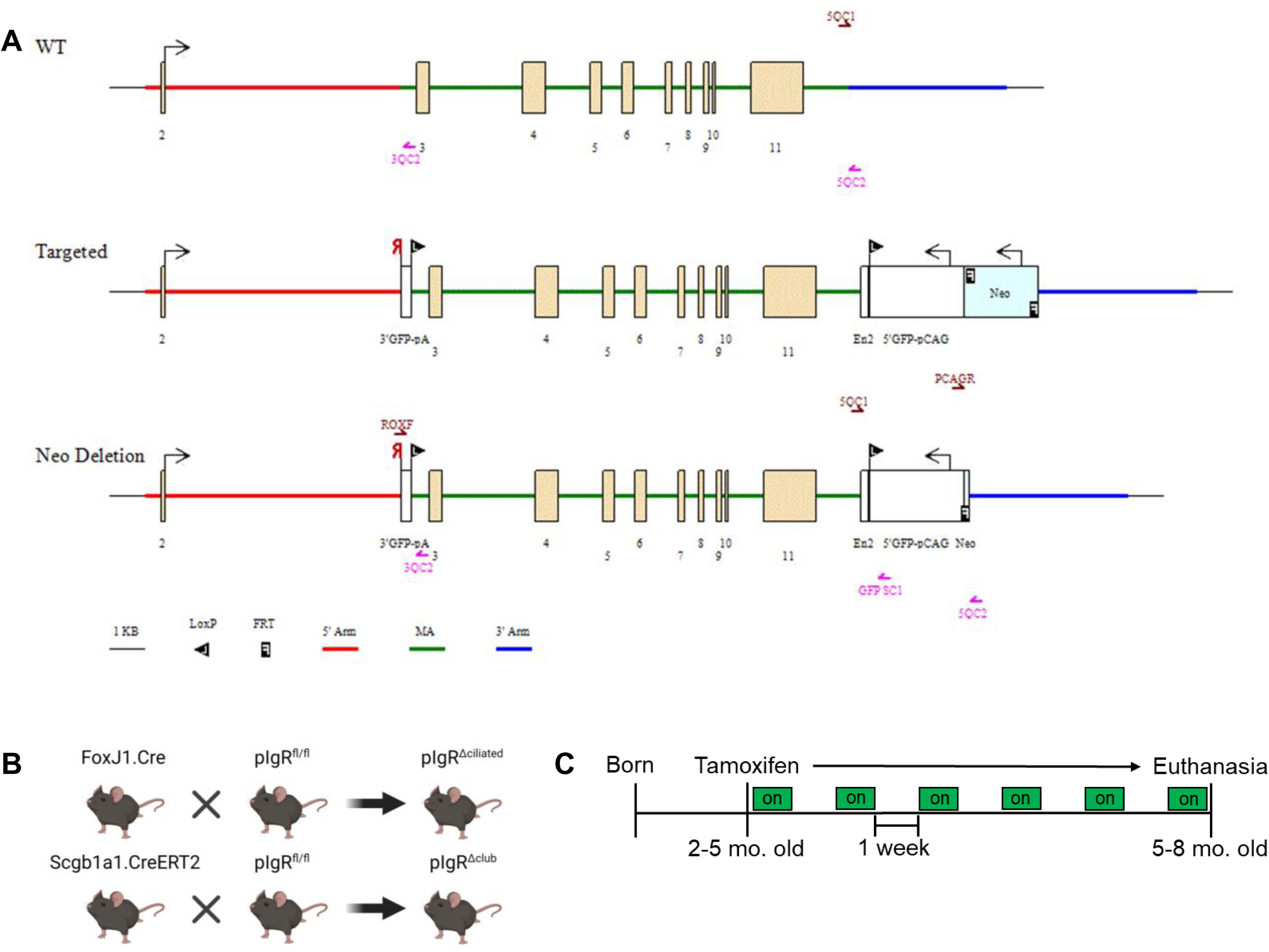
Construction of the pIgR^Δciliated^ and pIgR^Δsecretory^ mouse lines and tamoxifen administration schedule for pIgR^Δsecretory^ mice. (A) Design of the flox-pIgR mouse model. Schematic showing the *pIgR* gene region in wild-type C57BL6/J, flox-targeted, and neomycin-deleted flox-pIgR mice. LoxP sites flank exons 3 through 11 of *pIgR*. The coding regions for GFP are split between the upstream and downstream loxP sites such that Cre-mediated recombination results in GFP expression. FRT sites were used to remove the neomycin selection cassette; after neomycin deletion one 143 bp FRT site remains. The 5’ and 3’ regions outside of the targeting vector are shown in red and blue, respectively. (B) Diagram depicting generation of pIgR^Δsecretory^ and pIgR^Δsecretory^ mice by breeding flox-pIgR mice with FoxJ1.Cre and Scgb1a1.CreERT2 mice, respectively. (C) Schematic showing the timing of tamoxifen administration in pIgR^Δciliated^ mice. Tamoxifen chow was alternated with regular chow at weekly intervals to minimize toxicity.

**Supplemental Figure 4.**
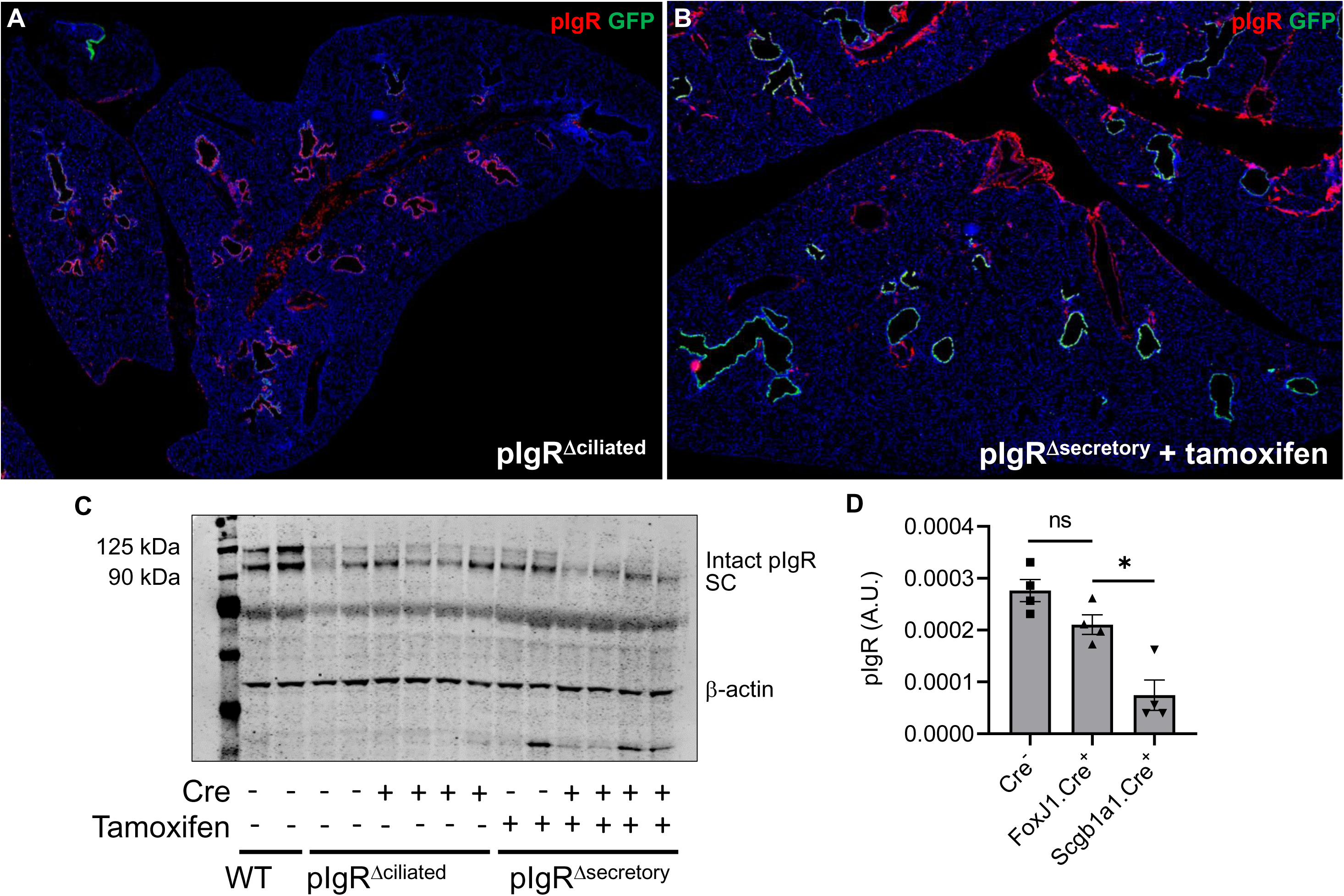
pIgR^Δsecretory^ mice have lower pIgR expression than pIgR^Δciliated^ or Cre^-^ littermate controls. (A, B) Representative low-power tiled images from an adult pIgR^Δciliated^ and pIgR^Δsecretory^ mouse showing pIgR and GFP staining in both lines. Identical acquisition settings were used for both images. (C) Immunoblotting for pIgR in whole-lung lysates from wild-type C57BL6/J, pIgR^Δciliated^, pIgR^Δsecretory^ mice and Cre^-^ littermate controls as indicated. β-actin is included as a loading control. (D) Densitometry-based measurement of pIgR expression in pIgR^Δciliated^ and pIgR^Δsecretory^ mice and Cre littermate controls.

**Supplemental Figure 5.**
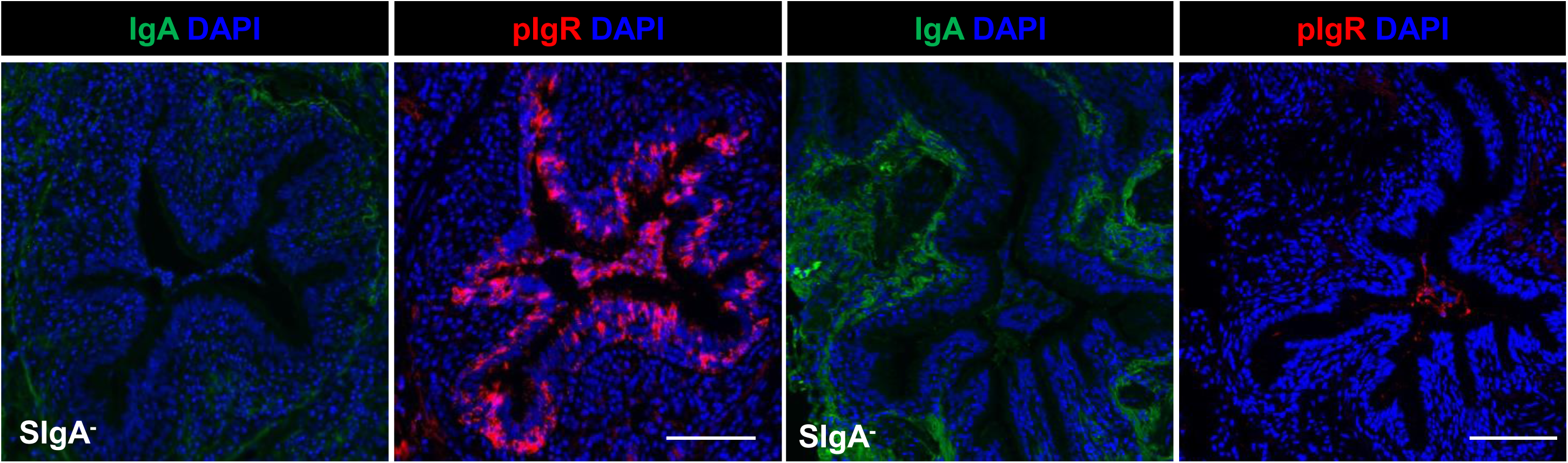
IgA and pIgR staining on serial sections of two different SIgA negative airways showing high (left) or low pIgR expression. Scale bars = 100 µm.

## Notes

### Competing Interest Statement

The authors have declared no competing interest.

