## Supplemental Table 1 for "Secretory cells are the primary source of pIgR in small airways"

**Supplemental Table 1. Demographic and clinical characteristics for human lungs.**

|  | Single cell RNA-sequencing | | RNA *in situ* hybridization | | RNA *in situ* hybridization / pIgR Immunofluorescence in SIgA- vs SIgA+ Airways |
| --- | --- | --- | --- | --- | --- |
|  | *Controls*  *n=10* | *COPD Patients*  *n=11* | *Controls*  *n = 6* | *COPD Patients*  *n=6* | *COPD patients*  *n=5* |
| Age  mean (range) | 28.6 (17-54)  2 Unknown | 60.3 (53-70) | 24 (9-41) | 57.7 (55-63) | 58.6 (56-63) |
| Race/ethnicity | NHW (6)  Unknown (4) | NHW (10), Black/AA (1) | NHW (5)  Unknown (1) | NHW (6) | NHW (4), Black/AA (1) |
| Sex | Female (1), Male (6)  Unknown (3) | Female (4), Male (7) | Female (4), Male (2) | Female (2), Male (4) | Female (1), Male (4) |
| Smoking history (pack-years)  mean (range) | 16.3 (1-43)  5 Unknown | 33.6 (5*-60)  3 Unknown | 6.2 (0-30) | 40 (20-60) | 46.4 (35-60) |
| Current smoker?  n(%) | 6 (60%)  3 Unknown | 0 (0%) | 3 (50%) | 0 (0%) | 0 (0%) |
| FEV1/FVC%  mean (range) | - | 31.5 (26-38)  3 Unknown | - | 30.8 (19-37) | 28.0 (21-34) |
| FEV1 (L,% pred)  mean (range) | - | 0.67, 21.8 (0.31-0.90, 13-30)  3 Unknown | - | 0.71, 21.8 (0.47-0.90, 13-28) | 0.70, 20.8 (0.31-1.02, 10-30) |
| FVC (L, % pred)  mean (range) | - | 2.15, 52.9 (0.97-2.97, 32-70)  3 Unknown | - | 2.36, 55.0 (1.55-2.88, 46-63) | 2.50, 57.0 (0.97-3.97, 32-91) |
| TLC (L, % pred)  mean (range) | - | 8.40, 141.5 (5.76-12.36, 107-177)  3 Unknown | - | 8.61, 132 (5.76-12.36, 107-177)  3 Unknown | 8.05, 131 (6.53-9.35, 114-157)  1 Unknown |
| DLCO (% pred)  mean (range) | - | 15.3 (7-30)  3 Unknown | - | 16.3 (7-30)  3 Unknown | 24 (10-49)  1 Unknown |

Abbreviations: NWH= Non-Hispanic White, AA= African-American. * indicates frequent marijuana use in addition to cigarettes.
